# Consequences of measurement error in qPCR telomere data: A simulation study

**DOI:** 10.1101/491944

**Authors:** Daniel Nettle, Luise Seeker, Dan Nussey, Hannah Froy, Melissa Bateson

## Abstract

The qPCR method provides an inexpensive, rapid method for estimating relative average telomere length across a set of biological samples. Like all laboratory methods, it involves some degree of measurement error. The estimation of relative telomere length is done subjecting the actual measurements made (the Cq values for telomere and a control gene) to non-linear transformations and combining them into a ratio. Here, we use computer simulations, supported by mathematical analysis, to explore how errors in measurement affect qPCR estimates of relative telomere length, both in cross-sectional and longitudinal data. We show that errors introduced at the level of Cq values are magnified when the TS ratio is calculated. If the errors at the Cq level are normally distributed and independent of telomere length, those in the TS ratio are positively skewed and proportional to telomere length. The repeatability of the TS ratio declines abruptly with increasing error in measurement of the telomere sequence and/or the control gene. In simulated longitudinal data, measurement error alone can produce a pattern of low correlation between successive measures of telomere length, coupled with a strong dependency of the rate of change on initial telomere length. Our results illustrate the importance of control of measurement error: a small increase in error in Cq values can have large consequences for the power and interpretability of qPCR estimates of relative telomere length. They also illustrate the importance of characterising the measurement error that exists in each dataset—coefficients of variation are generally unhelpful, and researchers should report standard deviations of Cq values and/or repeatabilities of TS ratios—and allowing for the known effects of measurement error when interpreting patterns of TS ratio change over time.

## Introduction

The length of telomeres—DNA-protein caps on the ends of linear chromosomes—has emerged across several fields as a key integrative biomarker to be studied in relation to ageing [1,2], environmental exposures [3], early-life experience [4,5], social determinants of health [6], stress [7], disease [8], and reproduction [9]. The widespread use of telomere length as a biomarker in epidemiological and ecological studies depends on the availability of a convenient and high-throughput method of estimating the relative average telomere lengths of a sample of individuals. That method comes from quantitative PCR (qPCR) [10]. In a recent large meta-analysis of the human telomere epidemiology literature, qPCR was used in 80% of the 143 studies, including almost all the studies with a sample size greater than 100 individuals [11]. The cheapness and simplicity of the qPCR method is a key enabling factor for the explosion of interest in the field of *in vivo* telomere dynamics.

There has been considerable debate concerning the impact of measurement error on the reliability of qPCR relative telomere length measurement. This debate concerns such issues as, for example, how much less reliable qPCR measurement is than other, more time-intensive methods [12–15]; what the sources are of variability in measurements [16–20]; and laboratory best practices for keeping measurement error to a minimum [21,22]. The purpose of this paper is rather different: regardless of what the source of measurement error is, what are its typical consequences for our datasets? Measurement error is classically modelled as the addition of a normally-distributed ‘noise’ term, whose standard deviation can be large or small depending on the precision of the technique, to the true value of the underlying quantities being measured. However, in qPCR telomere studies the actual laboratory values measured are first subjected to a non-linear transformation, and then combined into a ratio in order to estimate relative telomere length (the T/S ratio; henceforth we omit its ‘/’ to avoid confusion in formulae). Moreover, in longitudinal studies, the outcome variable is often the difference between two TS ratios. The likely consequences of measurement error for such variables as the TS ratio, or the change in TS ratio, are thus not obvious. We therefore sought to examine them through computer simulation of qPCR datasets, in which we could incorporate different amounts of error at the level of actual laboratory measurements, and examine the consequences of this for the outcome variables that qPCR telomere studies typically use.

The qPCR method of telomere length measurement follows the general principles of real-time DNA amplification using PCR: primers are used to amplify specific DNA sequences from a DNA sample; a fluorescent reporter allows detection of the abundance of the amplicon; and the measured variable is the Cq, the fractional number of PCR cycles required for a pre-chosen threshold of fluorescence to be reached. Because amplified DNA doubles successively during the PCR (assuming effectively perfect efficiency), Cq values should be linearly related to the base-two logarithm of the amount of the complementary sequence to the primer [10]. Thus, 2^−*Cq*^ is taken to be proportional to the amount of the target DNA sequence in the sample. (It is possible to incorporate imperfect amplification efficiency by using the measured slope of a standard curve [23] rather than 2, but that does not change the general principles that follow).

The amount of telomeric DNA present in a sample is the product of how many copies of the genome are present and the amount of telomeric DNA per genome copy. Hence, to estimate relative telomere length, it is important to normalize for the amount of DNA in the sample. This is done by amplifying a control genetic sequence that does not vary in copy number. Following Cawthon’s original terminology [10] we refer to this control sequence as the single-copy gene, although in fact all that matters is that its copy number is non-variable. The Cq for the single-copy gene is again transformed to 2^−*Cq*^. The critical estimator of relative telomere length—the TS ratio—has 2^−*Cq*^ for telomere in its numerator, and 2^−*Cq*^ for the single-copy gene in its denominator.

Our simulation approach is based on generating large datasets in which we first generate ‘true’ distributions of telomere length and of the number of genome copies in each sample. We then generate Cq values that reflect these quantities, but also incorporate random, normally-distributed measurement errors of varying magnitudes. We then use the Cq values to compute TS ratios, or the change in TS ratio for longitudinal cases. For simulations, unlike the usual empirical situation, we know what the ‘true’ underlying variables are, and thus we are able to compute the magnitude of the deviations between true and measured values, as well as other measures of reliability. Many of our key results were also derivable analytically, and these analytical findings are reported in the Appendix, section 1, and referred to in Results where relevant. Analytical and simulation findings were always concordant. In order to validate our assumptions, parameterize our simulations, and compare simulated to empirical outcomes, we also drew on two sets of empirical human qPCR data—one in which the same biological sample was measured multiple times under standard conditions, and a second that compared the results of multiple longitudinal human telomere studies.

## Methods

### Basic simulation framework

In all simulations, we first assign each biological individual in a cohort of *n* individuals a true average telomere length (*tl*). This is a normally-distributed quantity with mean 1 and specifiable standard deviation *σ_t_*. The variable *tl* represents how much longer or shorter than a typical individual that particular person’s telomeres are; thus, it is the true biological quantity that we wish to estimate by calculating a TS ratio from qPCR data.

Next, we generate DNA samples from each individual. The amount of single-copy-gene DNA in each sample, *DNA_s_*, is drawn from a normal distribution with mean *μ_s_* and standard deviation *σ_s_*.

The true amount of telomeric DNA in a given sample can thus be calculated:

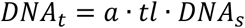

Here, *a* is a scaling constant (*a* ≫ 1) representing how many fold more abundant the telomeric sequence is than the single copy sequence in the average genome.

Now, we assume that qPCR is performed. In the ideal situation (no measurement error), since the *Cq* value from qPCR is linearly and negatively related to the base-2 logarithm of the amount of DNA in the sample, the error-free values of the *Cq* for the single-copy gene and for the telomeric sequence would be as follows (the *i* before the variable name indicates the ideal, error-free value):

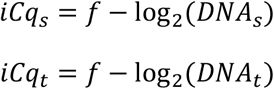

Here, *f* represents a constant set by the chosen fluorescence threshold.

Next, we introduce measurement error. We model this by the addition of a normally distributed measurement error term to each error-free Cq value. The Cqs that would actually be measured are thus as follows (the *m* indicates the measured as opposed to the ideal value):

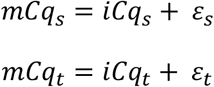

The measurement errors *ε* are drawn from normal distributions with mean 0 and standard deviations of *σ_εs_* and *σ_εt_* respectively. We henceforth refer to *σ_εs_* and *σ_εt_* as the ‘error σ’ for the single copy gene and telomere assay respectively. Our assumption unless otherwise stated is that the *ε_s_* and *ε_t_* are uncorrelated. However, there are circumstances in which this may not hold and the errors may be positively correlated; we explore the consequences of this in the Appendix, section 2.

The measured Cq values are combined to give the TS ratio. In its simplest form, this is given by:

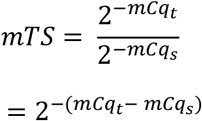

In practice, empiricists typically use Cqs measured from a standard sample to normalize their TS ratios. These standard Cq values can come from one individual sample, a pool of samples, or even the mean of the Cqs from all the samples. They are constants for any given study. As such, their effect is simply to rescale the TS ratio. Since the TS ratio is only a relative measure of telomere length, any constant rescaling has no impact on its reliability or relative precision. In our simulations we use the mean Cq for the telomere assay and single-copy gene in the whole cohort as the reference values. This has the effect of making the mean TS ratio approximately 1. No conclusions would be altered by using zeroes or two other values for the reference values. Incorporating the values from the standard samples (*rCqs*), the full formula for the TS ratio is:

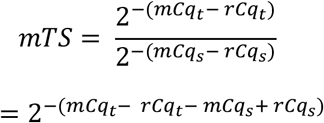

For each individual in all our simulations, we saved the true value of *tl*, alongside the measured Cqs (*mCq_t_* and *mCq_s_*) and TS ratio (*mTS*). We also saved the ideal values of the Cqs (i.e. the values that would have been observed had there been no measurement error; *iCq_t_* and *iCq_s_*). From these we could also calculate an ideal TS ratio (*iTS*). By subtracting the ideal from measured values, we were able to characterise the measurement error in each variable.

Default parameter values for the simulations were chosen in light of known telomere biology, and so as to produce values similar to those seen in empirical studies (table 1). Results were generally robust to numerical variation in the parameter values chosen. The code for the simulations is freely available at https://zenodo.org/record/1994387, and instructions for using it are included in the Appendix, section 3.

**Table 1.** Default values chosen for simulation parameters.

| Parameter | Description | Default value | Comment/justification |
| --- | --- | --- | --- |
| $n$ | Number of individuals in the cohort | 10000 | Interested in establishing patterns with high statistical power |
| $\mu_s$ and $\sigma_s$ | Mean and standard deviation across individuals of true amount of single-copy sequence present in sample | 10 and 1 | Variation in amount of DNA present in each sample assumed to be small relative to the mean amount |
| $a$ | Constant to represent how much more abundant the telomere sequence is than the single-copy sequence in the average genome | 1000 | Telomeric sequence many-fold more abundant than single copy gene in genome; Cq values for single copy gene typically more than double those for telomeric assay in real datasets [20]. |
| $\sigma_t$ | True standard deviation across individuals in relative abundance of the telomere sequence in the genome | 0.1 | Inter-individual standard deviation of adult telomere length measured by terminal restriction fragment is of the order of 10% of the mean (humans: 700bp / 7000bp [24,25]) |
| $f$ | Fluorescence threshold | 28 | Produces Cq values in similar range (around 10 to 25) to empirical data [18,20] |
| $\sigma_{\epsilon s}$ and $\sigma_{\epsilon t}$ | The error $\sigma$ for single-copy gene and telomere respectively; effectively, the standard deviation when the Cq of the same sample is measured many times | 0 to 0.3 | Examined range from 0 to well above values likely to be encountered in practice (which may typically be of the order of 0.05, see Results) |

### Simulation applications

Our initial investigations involved simulating datasets with different values of the error σs, to understand how measurement error in the Cqs affected the distribution of errors in *mTS*, and the relationships of *mTS* and its error to *tl*.

In addition to simulations of a single dataset, we performed simulations where sampled twice from the same *n* individuals, where the underlying telomere lengths *tl* were assumed to have not changed at all. A number of different analyses were possible using these repeated-sample datasets. We calculated the repeatability of *mTS*, that is, the extent to which it produces the same result when performed again on the same individuals in the absence of any true change. Repeatability can be assessed using the intra-class correlation coefficient (ICC) [26], here implemented as the ‘consistency’ ICC from the R package ‘irr’. We also used our repeated-sample simulated datasets as if they were the time 1 and time 2 measures from longitudinal studies where individuals’ true relative telomere lengths have not in fact changed during the study period. This allowed us to explore the consequences of varying the error σs for the apparent change in *mTS* between first and second measurement points (Δ*mTS*).

### Empirical datasets

We drew on two sets of empirical data, The first dataset (henceforth, dataset 1) comes from a recent methodological study using human samples [20] (we thank the study authors for generously providing us with their data in a rawer form than the published dataset). As part of this study, the same human DNA sample was run a total of 1728 times for telomere and 1728 times for single-copy gene over several plates, using three different light cyclers. This dataset is thus ideal for estimating the variability due to measurement error, since the biological sample, and hence relative telomere length, is always the same. We used only the light-cycler 1 data (576 telomere and 576 single-copy measurements over two sets of plates), since any normal study would be likely to analyse all samples on the same equipment.

The second dataset (dataset 2) comes from a recent paper on longitudinal studies of human telomeres [27]. Bateson, Eisenberg and Nettle collated data from seven human cohort studies in which telomere length had been measured twice in the same individuals, an average of 8.5 years apart (range 6.0 to 9.5 years; see [27] for full details). Five studies used qPCR, and two measured terminal restriction fragment using Southern blot. Of the various data collated, we extracted the correlation coefficient between the time 1 and time 2 telomere length measurement, and the correlation coefficient between the time 1 measurement and the change in telomere length between time 1 and time 2.

## Results

### Validating and parameterizing the simulation framework

All our simulations are based on the assumption that measurement error can be modelled as the addition of a normally-distributed noise term to the ideal Cq values for telomere and single-copy gene. In order to validate this assumption and gain a plausible range for the values of error σ in the simulations, we examined the distribution of Cq values in dataset 1 (which was, to recall, composed of the same biological sample run many times over two sets of plates). The observed Cq values did vary, and inspection suggests this variation can be reasonably modelled as the addition of a normal random variable to the central *Cq* value (figure 1). For telomere, the standard deviation of the Cq distribution was 0.053. This was not an artefact of combining data from two plates: for each plate separately, the standard deviations were 0.050 and 0.055. For the single copy gene, the standard deviation of the Cq distribution was 0.095. Again, there was variability within each plate (standard deviations per plate 0.091 and 0.098).

**Figure 1.**
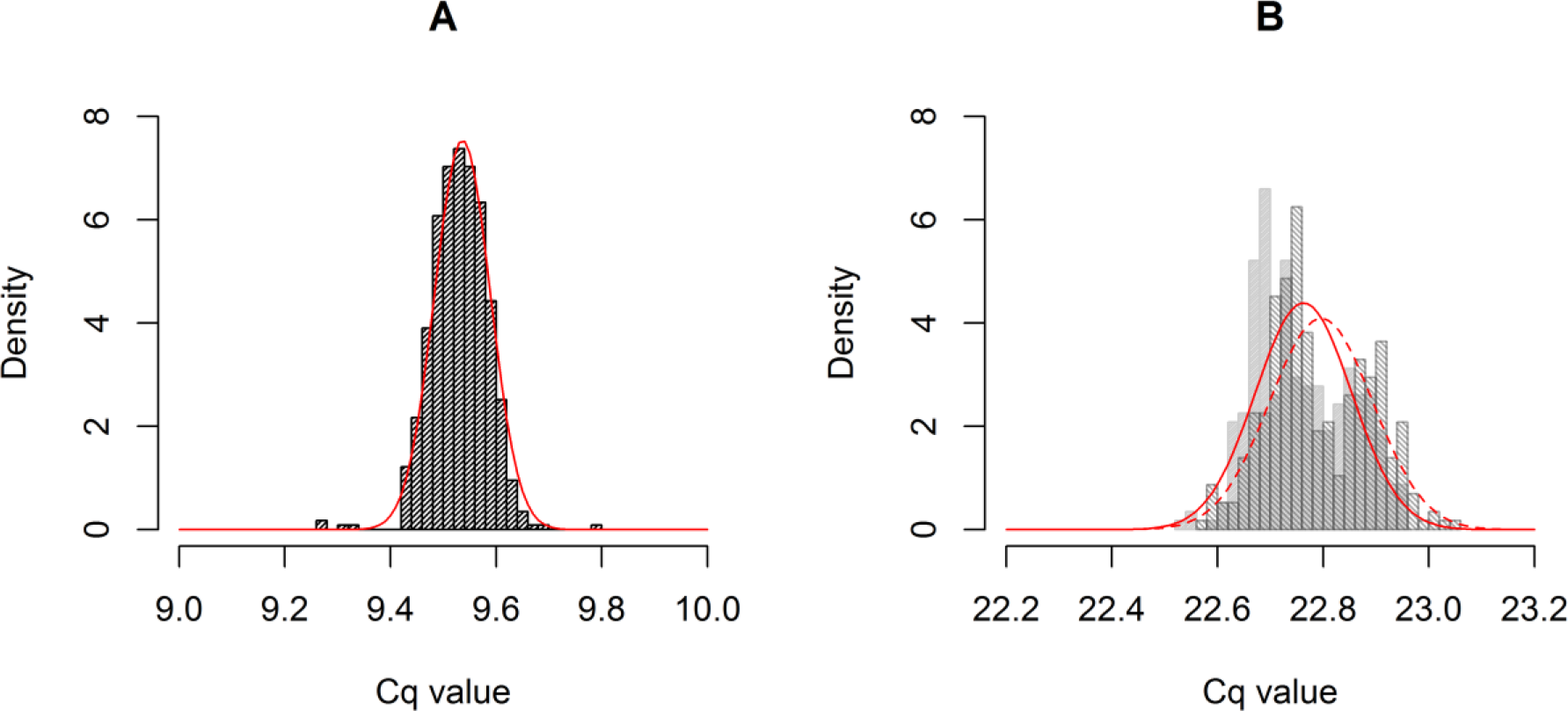
Observed distributions of Cq values for telomere (A) and single-copy gene (B) in dataset 1. For the single-copy gene where there the mean Cq varied significantly by plate, each plate is shown in a different shading. Lines show a superimposed normal density (separately by plate for single-copy gene).

In qPCR telomere studies, samples are usually run in triplicate and the replicate Cq values averaged to reduce measurement error. Assuming that adjacent wells are largely independent of one another, the consequence of this should be to reduce the effective error σ by a factor of √3 or approximately 1.73. Accordingly, in dataset 1, the standard deviation of means of three adjacent samples are 0.034 for telomere and 0.080 for telomere and single copy gene respectively (i.e. reductions by factors of 1.55 and 1.20 respectively compared to no replication). In the simulations that follow, the error σ discussed is the *effective* error σ after any averaging together of technical replicates has been carried out. The values from the dataset 1 suggest that post-averaging error σs of the order of 0.05 may be usual. Since our aim is to explore the potential consequences of increasing measurement error, we will consider error σ values from 0 all the way to 0.3.

### Consequences of error in Cq for TS ratio values

We simulated datasets with the error σs set to 0.05 for both reactions, and all other parameters at their default values. We calculated the error (measured minus ideal value) for each *mCq*, and also for each *mTS*. We scaled these by the standard deviation of the ideal Cq values and TS, so that ±1 indicates under- or over-estimating by a standard deviation of the ideal quantity under examination. Figure 2 plots the distribution of errors. As the figure shows, with these parameter values, the spread of relative errors in the *mTS* is larger than that in either of the *mCqs* (standard deviations for one run: A: 0.23; B: 0.37; C: 0.52). Whereas the errors in the *mCqs* are normally distributed (by assumption), the distribution of errors in *mTS* is not. For example, for one run of the simulation skewness values (with p-values from Agostino tests for skewness) were: error in *mCq* for telomere 0.005 (p = 0.83); error in *mCq* for single copy gene 0.003 (p = 0.90); error in *mTS* 0.12 (p < 0.001). Kurtosis values (with p-values from Shapiro-Wilks-Chen tests) were: error in *mCq* for telomere 0.01 (p = 0.92); error in *mCq* for single copy gene -0.05 (p = 0.32); error in *mTS* 0.19 (p < 0.001). Thus, even if the errors introduced in measuring the Cq values are normally distributed, the error in the computed TS ratio is both positively skewed (more large overestimates than large underestimates), and leptokurtic (more extreme outliers than would be found in a normal distribution). In the Appendix (section 1) we show that this is because the error in the TS ratio belongs to a class of distribution known as a normal-log-normal mixture distribution; these distributions are generally skewed and leptokurtic [28].

**Figure 2.**
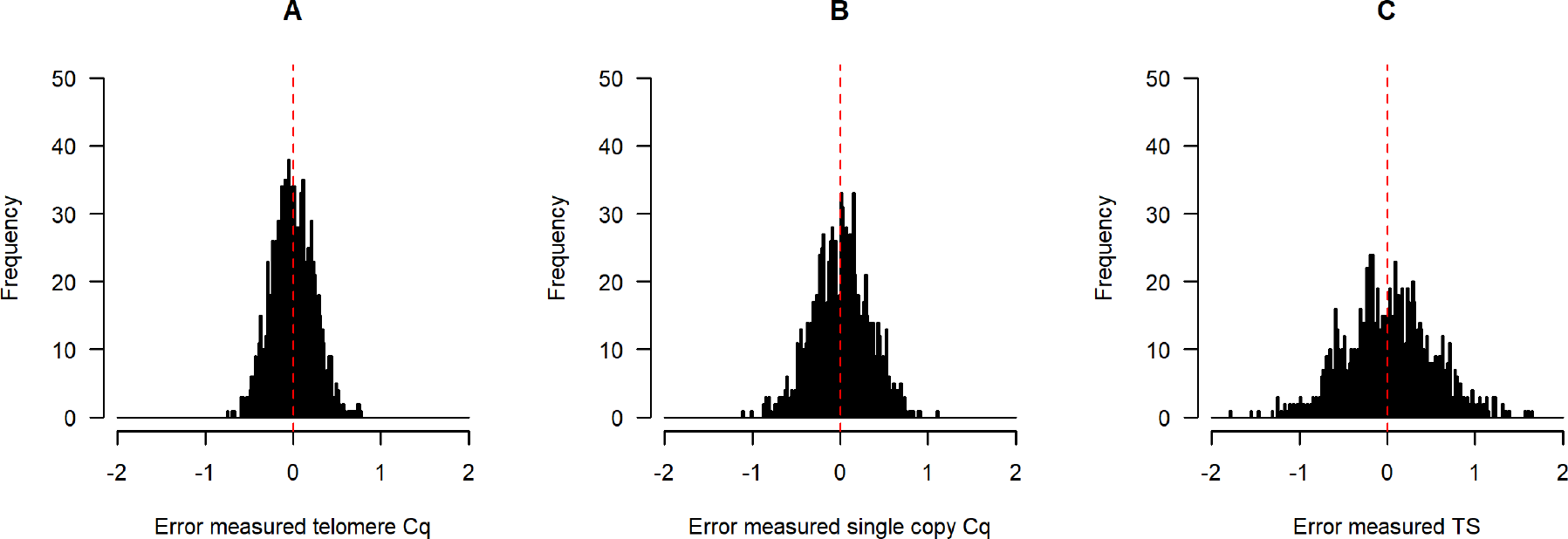
Distribution of errors in Cq values (telomere, A, and single-copy gene, B) and the resulting TS ratio (C) for runs of the simulation with *n* = 1000 and *σ_εs_* = *σ_εt_* = 0.05. The scale in each case is standard deviations of the ideal (error-free) quantity in the simulated sample.

### Relationship between measured TS ratio and telomere length

We examined the association between *tl* and *mTS* for simulated datasets with three different levels of error σ (figure 3). With *σ_εS_ = σ_εt_* = 0, *mTS* is perfectly correlated with *tl*, as expected (see also Appendix, section 1, results 1 and 2). As the error σs increase, there is increasing scatter in the association of *mTS* to *tl* (figure 3, panels B, C), and the scatter is greater for longer telomere lengths (figure 3, panels E, F). We confirmed analytically that the expected magnitude of the measurement error in *mTS* is proportional to *tl* and hence greater for individuals with longer telomeres (see Appendix, section 1, result 3).

**Figure 3.**
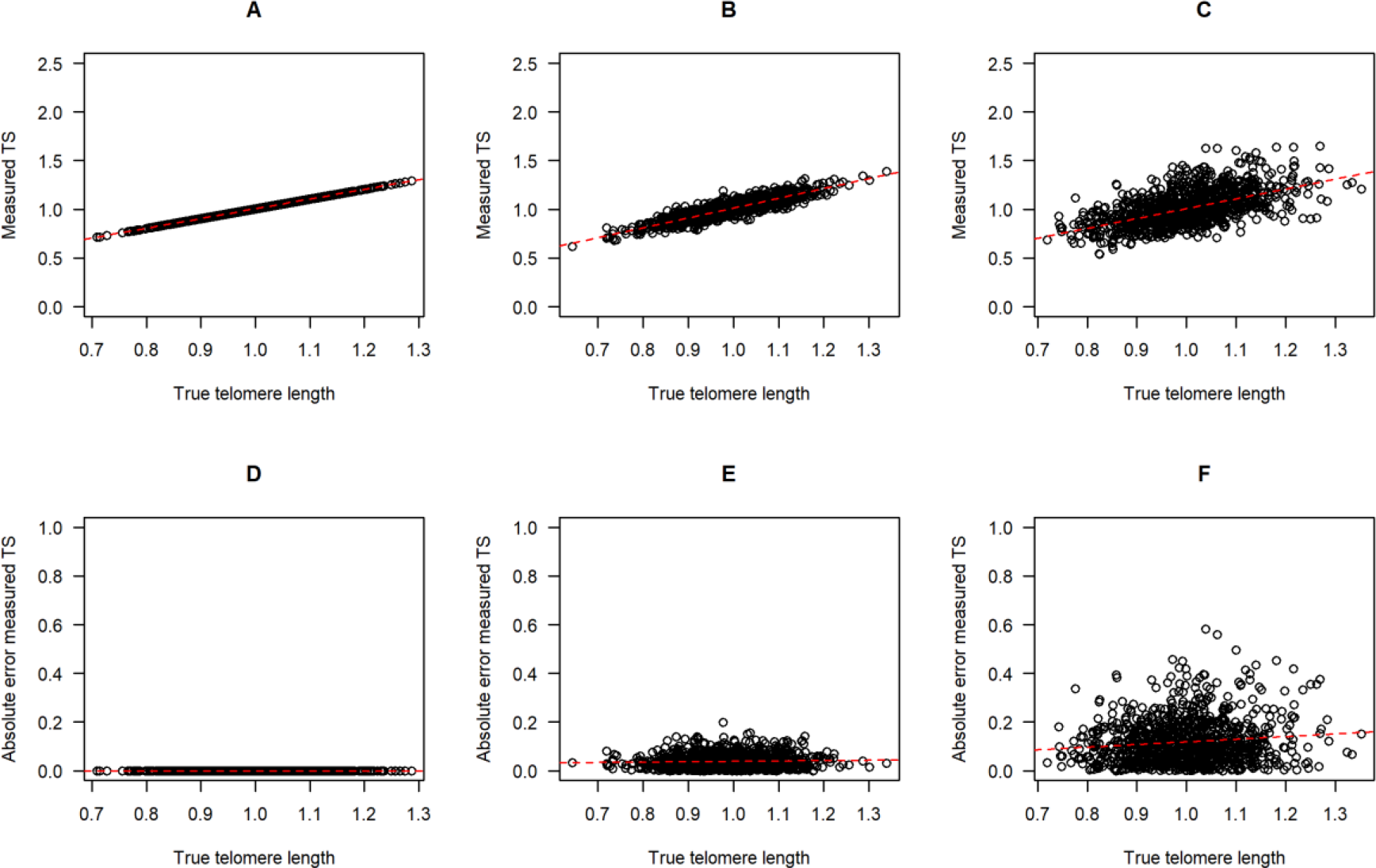
Association between measured TS ratio and true telomere length as measurement error increases. Top row: scatterplots of *mTS* against *tl* for *σ_εs_ = σ_εt_* = 0.00 (A); *σ_εs_ = σ_εt_* = 0.05 (B); and *σ_εs_ = σ_εt_* = 0.15 (C). Bottom row: The absolute magnitude of the difference between *mTS* and *iTS*, for *σ_εs_ = σ_εt_* = 0.00 (D); *σ_εs_ = σ_εt_* = 0.05 (E); and *σ_εs_ = σ_εt_* = 0.15 (F). Red dotted lines indicate linear regressions of *mTS* on *tl*.

### Repeatability of the measured TS ratio

We next examined the repeatability of *mTS* by simulating datasets where two separate samples are taken from each biological individual, and true *tl* is unchanged. The repeatability (intra-class correlation) should therefore be equal to 1, and any deviation from 1 reflects measurement error. figure 4A shows how repeatability varies with *σ_εs_* and *σ_εt_*. Thus, figure 4A suggests that error σs of less than around 0.08 are required for repeatability of greater than 0.75 in *mTS*; and that error σs of greater than around 0.11 will produce *mTS* whose repeatability is less than 0.6. The two error σs affect repeatability equally and symmetrically. An alternative to calculating repeatability for these datasets would be to calculate the correlation coefficients between either of the *mTS* and the *tl*. Such a calculation gives a very similar pattern to figure 4A. Indeed, the repeatability of *mTS* and the correlation between *mTS* and *tl* are closely linked: when repeatability is high, it is because both *mTS* values are highly correlated with *tl*, and hence with one another.

The advantage of calculating *mTS* over just using the raw telomere Cq (or 2^−*Cq*^) as the estimator of relative telomere length is that it corrects for variation in the amount of DNA present. However, calculating *mTS* also has the drawback of introducing a second source of measurement error. Figure 4B fixes the telomere error σ at 0.05 and examines how varying the error σ for the single-copy gene affects the repeatability advantage of calculating *mTS* over the *mCq_t_*. *mTS* is much more repeatable than *mCq_t_* (and also much better correlated with *tl*) when *σ_εs_* is small, but the gap reduces sharply as *σ_εs_* increases. If *σ_εs_* reaches around 0.15 or more (under the parameter values used here), then *mTS* is no more repeatable than *mCq_t_*, since its advantage in controlling for DNA variation is entirely offset by the extra measurement error introduced by considering the single copy gene.

**Figure 4.**
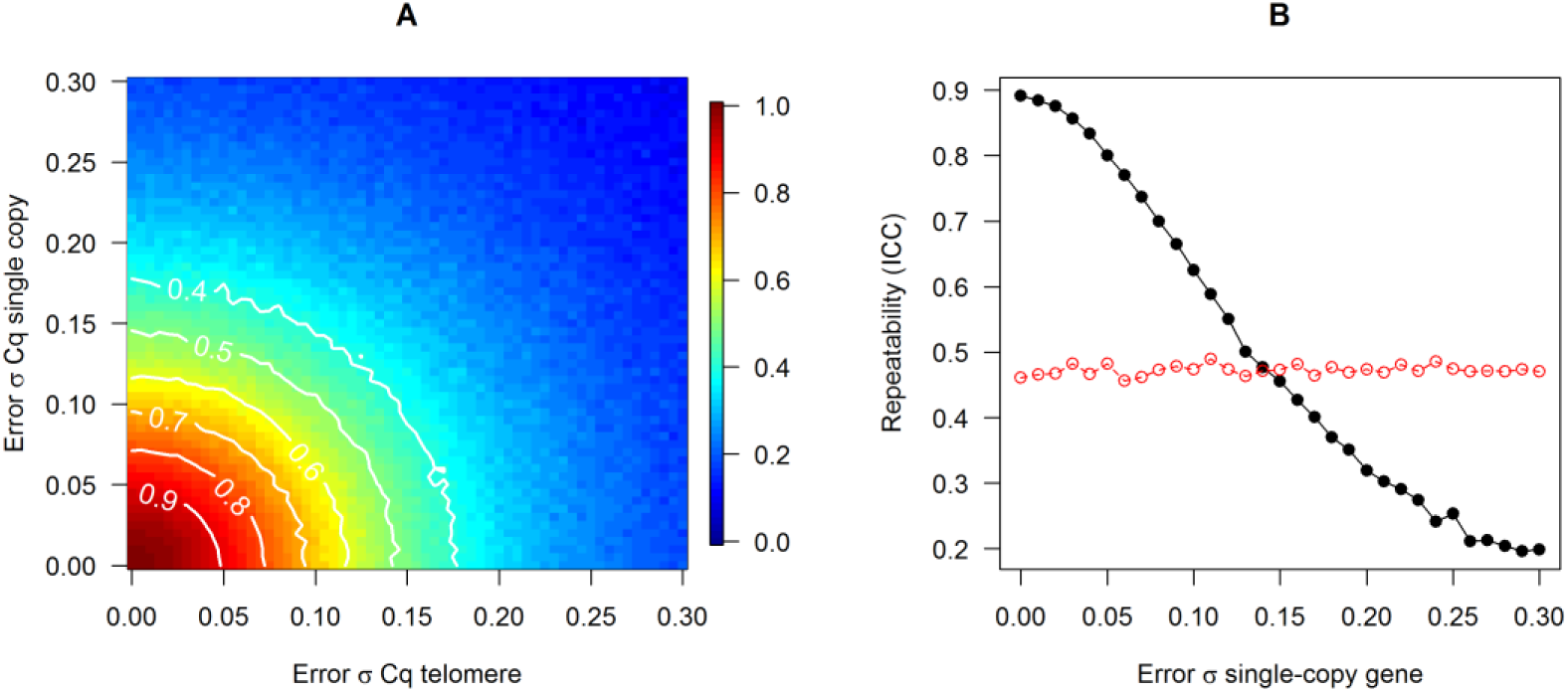
Effects of measurement error on repeatability of the measured TS ratio. A. Repeatability (intra-class correlation coefficient) of the TS ratio as measurement error in the two Cq values varies. B. Repeatability of the measured TS ratio (black line, filled circles) and the measured telomere Cq (red circles), as measurement error in the single-copy gene increases. The error σ for telomere is fixed at 0.05. The point where the two lines cross is the point where the advantage of controlling for sample-to-sample variation in the amount of DNA present is offset by the extra measurement error introduced.

We also investigated how the pattern of repeatability in figure 4A is affected by allowing the errors in the single-copy gene and the telomere reaction for the same biological sample to be non-independent. In general, positive correlation between the errors reduces the impact of measurement error in the Cqs on the TS ratio and its repeatability (see Appendix, section 2, figure S1). To see why this is the case, consider what would happen if the two errors were perfectly correlated: the measurement error in the telomere Cq would be matched by an identical error in the single-copy gene, the two errors would cancel, and the resulting TS ratio would be error-free (Appendix, section 1, result 5). However, the impact of more modest correlations on measurement error in the TS ratio is small.

### Association between successive measurements and regression to the mean

We again simulated datasets where the same biological individuals are measured twice with no true telomere length change. Figure 5 plots the association between *mTS* at the first time point and *mTS* at the second, for no measurement error (panel A), *σ_εs_ = σ_εt_* = 0.05 (panel B), and *σ_εs_ = σ_εt_* = 0.15 (panel C). As the error σs increase, the regression line through the data is rotated around the bivariate means and increasingly flattened relative to the line *y = x*. Thus, the variance in the second measurement explained by the first measurement (the r^2^, which in the absence of measurement error should be 1) declines (figure 5D). We also calculated the strength of association between *mTS* at time 1 and subsequent apparent change (*ΔmTS*) as error σs increase (figure 5E). With increasing measurement error, *ΔmTS* comes to depend increasingly strongly on time 1 *mTS*. This is a known effect due to regression to the mean: in two largely uncorrelated measurements, if the first is far from the centre of the distribution, the second will on average tend to be closer to the centre.

Figure 5F replots the simulated data from figure 5D and 5E so that the correlation in *mTS* between time 1 and time 2 is shown on the horizontal axis, and the correlation between *ΔmTS* and time 1 *mTS* is shown on the horizontal axis. The simulations suggest that a signature of measurement error in longitudinal datasets, if in fact true telomere length is largely stable, is the combination of low correlation between time 1 and time 2 *mTS*, and strong dependency of change in *mTS* on time 1 *mTS*. We would thus predict that where studies have a low time 1-time 2 correlation due to measurement error, they should also have a strong dependency of apparent telomere length change on time 1 telomere length. Data relevant to this prediction come from dataset 2, which reported the correlation between time 1 and follow-up TS ratio, and between time 1 TS ratio and change in TS ratio, for seven human cohorts in which relative telomere length had been measured twice. The empirical data are superimposed on figure 5F. Those studies that have a high correlation between time 1 and time 2 TS ratio (which were non-qPCR studies) show a negligible dependency of TS ratio change on time 1 TS ratio, whereas those with a low correlation between time 1 and time 2 show a strong dependency. Thus, one interpretation of these data is that some of the qPCR studies feature a high degree of measurement error, and show the predicted combination of low time 1-time 2 correlation and apparent dependence of the rate of change on the time 1 telomere length.

**Figure 5.**
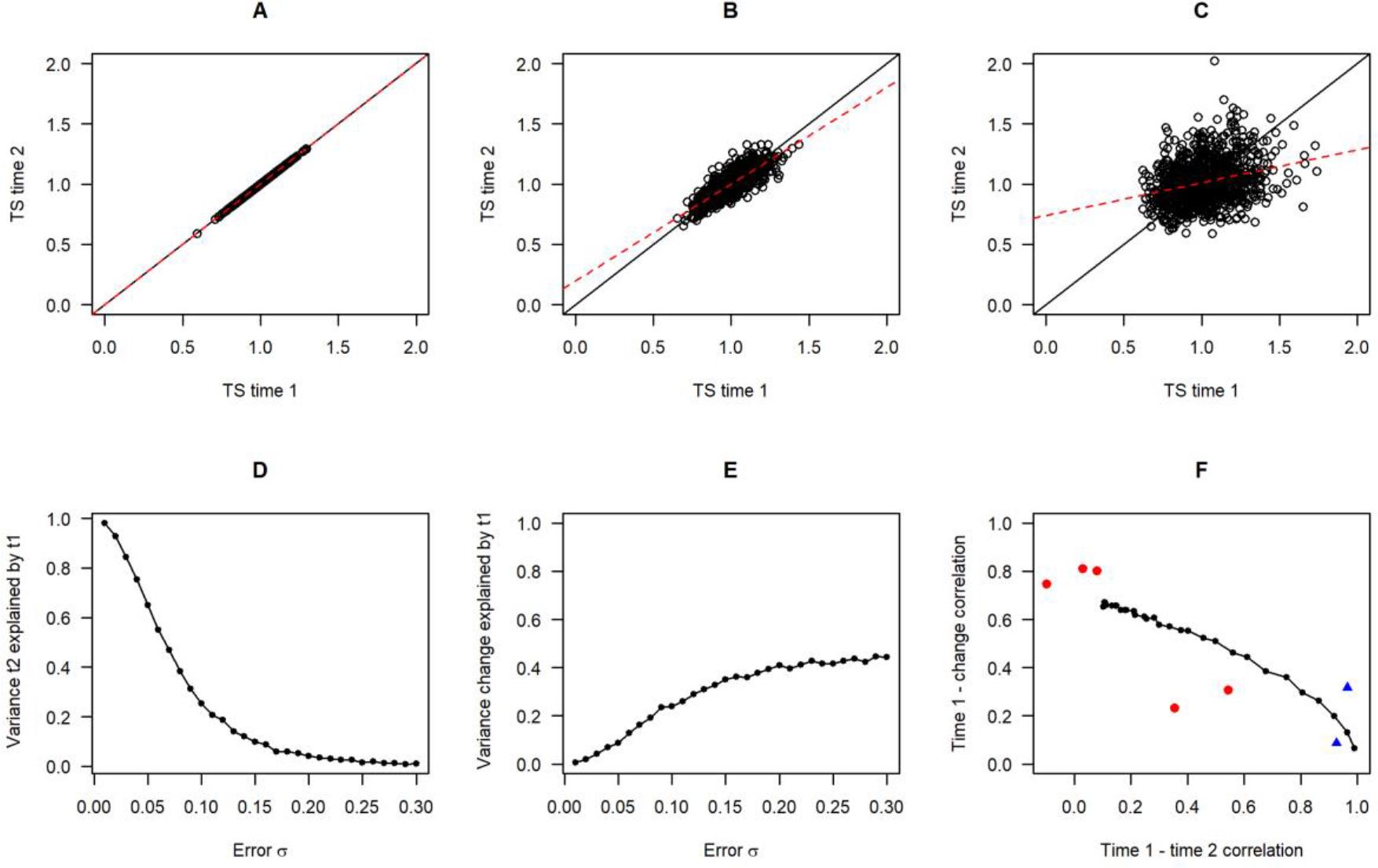
Consequences of measurement error for patterns in longitudinal datasets. Panels A to C: Association between *mTS* at time 1 and *mTS* at time 2 assuming no true change, for no measurement error (A), *σ_εs_ = σ_εt_* = 0.05 (B); and *σ_εs_ = σ_εt_* = 0.15 (C). Panel D: The variance in *mTS* at time 2 explained by *mTS* at time 1 with no true change and increasing levels of measurement error (*σ_εs_ = σ_εt_)*. Panel E: The variance in change in *mTS* between time 1 and time 2 explained *mTS* at time 1, assuming no true change and increasing levels of measurement error (*σ_εs_ = σ_εt_*). Panel F: Time 1-time 2 correlation against Change-time 1 correlation for simulated datasets (black circles and lines). Measurement error increases from 0 at bottom right to 0.2 at top left (*σ_εs_ = σ_εt_*). Superimposed are empirical values from the seven large human longitudinal cohorts from dataset 2. qPCR studies are shown in red and studies measuring terminal restriction fragment by Southern blot in blue.

## Discussion

Using a combination of computer simulation and mathematical analysis, we were able to elucidate some important features of the potential impact of measurement error in datasets where relative telomere length is estimated by calculating a TS ratio from qPCR. First, because of the way two independent measurement errors (in the telomere and single-copy gene reaction) are exponentiated and combined, any error at the level of Cqs is magnified into a proportionately larger error in the TS ratio. Confirming this, papers reporting some estimate of measurement error for both the individual Cqs and the TS ratio do report proportionately greater error for the TS ratio (e.g. [4,20]). Repeatability of the TS ratio is high (greater than 0.75) as long as measurement errors in Cq are of the order of 0.075 or less, but it declines rapidly as error in the Cqs becomes greater. To illustrate with some concrete numbers, according to our simulations, repeatability of the TS ratio should be about 0.80 with error σ values of 0.05, 0.51 with error σ values of 0.1, and 0.28 with error σ values of 0.15. A widespread conclusion when surveying the qPCR telomere epidemiology literature is that there is a great deal of heterogeneity between studies [29,30]. Our findings suggest that small differentials in errors in the laboratory would be sufficient to drive large heterogeneity in outcomes.

The general consequence of measurement error is to attenuate power to detect true associations. If measurement error in the TS ratio is substantial, the inferential security of resulting claims is undermined. Most obviously, the greater the measurement error, the more likely it is that reported null associations represent false negatives. Perhaps less obviously, ‘significant’ results that are found are more likely to represent false positives when measurement error is greater [31]. This is because, as measurement error increases and power declines, the rate of true positive findings reduces, but the rate of false positive findings remains the same (1/20 for a threshold *p* < 0.05). Thus, in the set of associations with *p* < 0.05, the ratio of true to false positives becomes worse.

The advantage of calculating a TS ratio, namely control for variation in DNA concentration in the sample, is substantial when the single-copy gene can be measured with low error, but eroded as measurement error in the single-copy gene increases. The simulations show that there is a level of single-copy gene measurement error at which the TS ratio becomes no more repeatable than the telomere Cq. The measurement error could well be worse for single-copy gene than for telomere: the precision of qPCR is thought to increase with copy number [32], and this is necessarily lower for the single-copy gene. In line with this, in dataset 1 analysed here, the error variation in Cq for single-copy gene was larger than for telomere. We are not suggesting that single-copy gene measurement error is typically large enough in practice to undermine the utility of calculating a TS ratio. However, according to our simulations, the measurement error in the single-copy gene would only have to be around twice what we observed empirically in dataset 1 for the TS ratio to be no more repeatable than the uncorrected Cq for telomere (under our simulation assumptions about the amount of true variation in DNA abundance). This is a rather clear illustration of why even modest increases in measurement error are corrosive in qPCR telomere studies.

Our main results are derived on the assumption that the errors in the Cqs of telomere and the single-copy gene are independent for a given biological sample. This may be a reasonable approximation, particularly for traditional methods where the two reactions occur in different wells. However, plate or well location effects, or issues with DNA extraction or purity, could affect both telomere and single-copy gene reactions for the same sample, and thus some correlation in errors cannot be dismissed as a possibility. Moreover, in the multiplex assay [33], the two reactions occur in the same well, and thus the scope for non-independence of the two measurement errors is even greater. The general consequence of non-independence is to reduce the impact of the measurement error at the level of the TS ratio. Thus, if the multiplex assay produces more highly correlated measurement errors, this is an advantage. In a sense, this advantage was already understood in the development of the assay: a key argument for it was to make sources of variability like pipetting affect telomere and single-copy gene alike [33]. Overall, though, our simulations show that modest non-independence between the two errors has only a very small mitigating effect on the consequences of measurement error for the TS ratio.

The error in the TS ratio is proportional to true telomere length, being larger for individuals with relatively longer telomeres. This is true even though the simulations assume that the measurement errors at the Cq level are independent of telomere length: it follows directly from the TS ratio formula. This is an issue with potentially complex consequences. For example, telomere length shortens rapidly with age in very young individuals [4,34]. In a cohort whose ages span this period, relative telomere length would be estimated with greater error in the youngest age group (equally, in an experimental design where the treatment has a dramatic effect on relative telomere length, the experimental group might be estimated with less or more error than the control group). This violates assumptions of homoscedasticity central to many statistical analyses. More generally, researchers often report that the distribution of TS they observe is positively skewed, and resort to logarithmic transformations to correct this (e.g. [35–38]). Our simulations suggest that measurement error will predictably produce this skew, and also considerable kurtosis (the presence of more extreme outliers than found in a normal distribution), even if the underlying distribution of relative telomere lengths is normal.

When applied to longitudinal studies, measurement error alone can produce a pattern of low correlation between the first and second telomere length measurements, coupled with a strong dependence of the apparent telomere length change on the initial telomere length. This pattern has already been recognized and discussed specifically in relation to telomere length [39–42], and in longitudinal data more generally [43]. It is not a consequence of the TS ratio in particular, but of any set of repeated measurements where there is measurement error. An implication is that because at least part of the apparent association of initial telomere length and subsequent change is spuriously created by measurement error, controlling for initial telomere length in regression models in which the outcome variable is telomere length change is often invalid and biases inferences [27]. Specifically, it biases the estimate of the effect on telomere length *change* of any predictor variable that is associated with telomere *length* at baseline.

Our longitudinal simulations are all based on the assumption that true telomere length is a highly stable individual characteristic over time. High individual stability, across adulthood at least, is what is seen in human longitudinal studies that measure telomere length with Southern blot ([44], see also figure 5F), assumed to be a higher-fidelity method than qPCR. If we assume that these studies capture a gold standard of what human telomere dynamics through adulthood are typically like, then the combination of low time 1-time 2 correlation and high dependency of change on time 1 length found in some human qPCR cohort studies (as shown here in figure 5F) probably suggests that these studies are characterised by a level of measurement error that seriously undermines reliability. However, we should be wary of inferring that just because measurement error alone can produce an apparently low correlation between time 1 and time 2 telomere length, then all such low correlations are necessarily attributable to measurement error. In populations living under variable ecological conditions, telomere length may truly be more dynamic over the course of life [3,45]. Likewise, it would be invalid to assume that because measurement error alone can produce an apparent association between initial telomere length and subsequent change, then all such associations are completely reducible to measurement error. On the contrary, there is evidence suggesting that longer telomeres may shorten faster even after correction for measurement error [39,41]. Such cases illustrate the importance of researchers understanding and characterizing the measurement error in their data. If the level of measurement error is known, then it is possible to generate the appropriate null hypotheses about what the association between initial and follow-up length, or between initial length and change, should look like if there are no biological dynamics at work. Our simulations allow for estimation of what the time 1-time 2 correlation should be in the TS ratio under the null hypothesis of no telomere length change, as long as the error σ values can be estimated from the data. For dependence of change on the initial length, our simulations or Blomquist’s formula (see [43]) provide simple ways of predicting what the apparent dependence should be under the null hypothesis if there is a specified level of measurement error.

The empirical value that comes closest to the error σ of our simulations is simply the standard deviation of Cq when replicates of the same samples are run repeatedly. This is one of two recommended reporting alternatives given in the MIQE guidelines for characterising measurement error [32]. These values can be compared directly to those in our simulation, recalling that averaging *k* technical replicates together reduces the effective error σ by √*k*, as long as we can assume that the replicates are independent.

In practice, more researchers appear to use the MIQE guidelines’ [32] second alternative reporting item, namely a coefficient of variation (CV). Though the guidelines prescribe a CV on the 2^−*Cq*^ values, and our experience is that it is often on the raw Cqs that a CV is reported; this has long been known to imply a misleadingly low level of measurement error [46]. The CV is problematic for a number of reasons, particularly when attempting to compare across studies or methods, as has been discussed elsewhere [13,29]. It can also give a misleading impression when comparing measurement error in telomere and single-copy gene: since the single-copy gene is much rarer than the telomere sequence, the denominator of its CV will be much larger (for Cq), or smaller (for 2^−*Cq*^). Thus, reporting CVs makes it non-obvious whether the measurement error for the telomere reaction is larger, smaller, or the same as those for the single-copy gene in absolute terms. Thus, we would strongly recommend reporting standard deviations of Cq from technical replicates in preference to CVs. In addition, the intra-class correlation coefficient of TS ratios from a suitable set of test samples run multiple times is an easily comprehensible summary of the repeatability of a measurement technique [13]. Our figure 4A shows that the value of intra-class correlation coefficient is directly governed by the effective error σs of the Cqs.

Our simulations have a number of limitations. First, we assume that each individual has a unitary relative telomere length. This is itself a simplification: every individual has a distribution of telomere lengths, and this distribution will vary between as well as within cells. Our simulations do not model this level of variability, but start from the assumption that relative telomere length can be adequately represented as a single quantity (an assumption central to the qPCR approach). Sample to sample biological variability (i.e. sampling or temporal variation in cellular composition of samples) is simply incorporated here with other sources of measurement error, rather than modelled as a biological process that may be of interest in its own right. Second, all of the measurement error we model is non-systematic, as exemplified by our use of independent drawings from a normal distribution of errors. In empirical studies, some error is however likely to be systematic, with consistent differences between plates, or well positions on a plate [17]. Where error is systematic, it can be mitigated through appropriate statistical correction. By carefully examining plate and well differences and correcting for them, researchers can thus reduce their effective error σ values somewhat further. Third, our simulations did not include any correction for amplification efficiency, although such corrections have been proposed and are often employed [23]. Our simulations effectively capture the case where amplification efficiencies are the same for telomere and the single copy gene (in which case amplification efficiency-corrected and uncorrected T/S ratio coincide). If this is not the case, correction may be applied. Thus, researchers have multiple ways of keeping the *effective* error σ values to a minimum; our simulations deal with the effects of the residual error that remains once all such steps have been taken. However, if statistical corrections include estimating parameters empirically (such as, for example, the estimating the amplification efficiency of each particular reaction), then of course more error is potentially introduced in the estimation, and that error gets incorporated into the statistical correction. Thus, the impact on the overall reliability of the relative telomere length measurement is hard to predict.

## Conclusions

We have presented a simple simulation framework for exploring the impact of errors in measurement of Cq values on estimation of relative telomere length measurement using the T/S ratio. The results illustrate the potentially large consequences for reliability of small increments in measurement error, and hence underline the need for researchers to both minimise and understand the measurement error that exists in their datasets. They also illustrate the value of simulation and mathematical analysis as tools for to guide empirical practices.

## Acknowledgements

We thank Sharon Savage and Casey Dagnall for their assistance with access to dataset 1.

## Appendix Section 1: Analytical treatment of measurement error in the TS ratio

This section examines the TS ratio under measurement error, providing analytical results to support simulation findings.

As specified in the main paper, the ideal (i.e. error-free) Cq values for the telomere assay and single copy gene relate to the amounts of each kind of DNA present in the sample as given in (1) and (2).

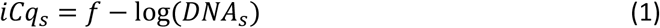

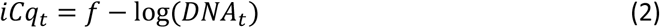

Here, *f* denotes a constant set by the chosen fluorescence threshold. The amount of telomeric DNA present is proportional to the amount of single copy gene DNA present, but scaled by the relative telomere length of the individual.

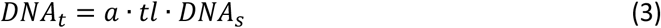

Hence, combining equations (2) and (3):

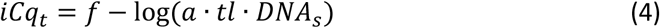

The measured Cq values are the true Cq values plus a measurement error term, as follows:

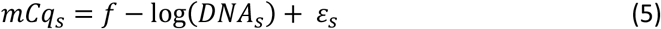

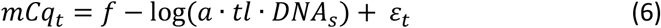

Here, *ε_t_* ~ *N*(0, *σ_εt_*) and *ε_s_* ~ *N*(0, *σ_εS_*). The formulae for the ideal and measured TS ratio are given in (7) and (8).

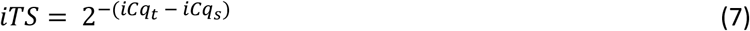

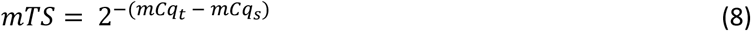

Reference Cq values for a standard sample are typically subtracted from the Cqs for the single copy gene and telomeric assay in calculating TS ratios. The effect of this is simply to rescale the TS ratio; such rescaling can be ignored in what follows without loss of generality, and hence for clarity we do not include this step here (though see main paper for the TS formula with these reference values included).

By substituting into (7) and (8) and rearranging, we have:

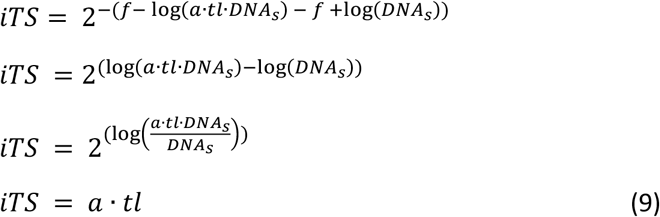

Thus, (9) gives us **Result 1:** The TS ratio, if measured without error, is proportional to the relative telomere length in the sample.

For the measured TS ratio where there is measurement error, we have:

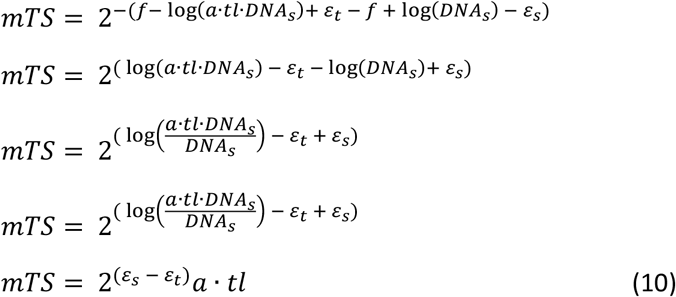

From (10), we have **Result 2:** The measured TS ratio is proportional to relative telomere length multiplied by 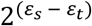, or two to the power of the difference between the measurement errors in the two Cq values.

The error in the measured TS ratio is the difference between *mTS* and *iTS*. From (9) and (10):

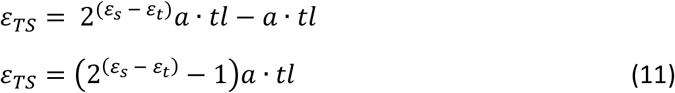

By inspection of (11), we have **Result 3:** The error in the TS ratio is proportional to telomere length. This is true even though the errors in the Cq values were assumed to be independent of the amounts of telomere and single copy DNA in the samples.

If *ε_t_* ~ *N*(0, *σ_εt_*) and *ε_s_* ~ *N*(0, *σ_εs_*), from properties of the normal distribution:

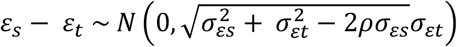

Here, *ρ* is the correlation between *ε_s_* and *ε*_*t*_. Hence, the distribution of *ε_TS_* is the distribution of:

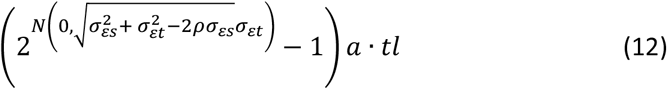

From (12), we can make the following inferences for the case where the measurement errors in the Cq values are normally distributed:

- **Result 4:** Positive correlations between *ε_s_* and *ε_t_* reduce the size of measurement errors in the TS ratio. From (12), given that *2σ_εt_σ_εs_* is positive, increasing *ρ* will always reduce the size of 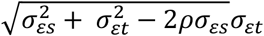, and hence the standard deviation of *ε_TS_*.
- **Result 5:** Perfect positive correlation between the measurement errors of the Cq for telomere and the Cq for the single copy gene eliminates measurement error in the TS ratio entirely, as long as the extent of measurement error is the same for the two reactions. Where *ρ* = 1 and 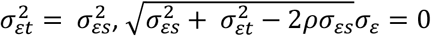. Hence, from (12), the measurement errors in the TS ratio are: (2^*N*(0,0)^ − 1)*a* · *tl* = 0.

If we can assume that telomere length itself is normally distributed, then we can see from (12) that the error in the TS ratio contains a normally distributed component (*tl*) and a log-normally distributed component (since the logarithm of 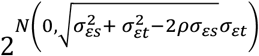 is by definition normally distributed, 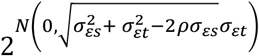 is log-normal). Thus, the distribution of *ε_TS_* belongs to the class of normal-log-normal mixture distributions. Such distributions are typically skewed and leptokurtic (Yang 2008).

## Section 2: Simulation results with correlations between errors

Simulation results reported in the main paper assume that the error in the telomere Cq and the error in the single-copy gene Cq are independent; that is, in the notation of section 1, *ρ* = 0. We repeated the main simulations assuming positive values of *ρ*. Increasing values of *ρ* attenuate the impact of measurement error at the Cq level on the TS ratio (see section 1, result 4). Although *ρ* = 1 makes the TS ratio error-free regardless of the magnitude of error in Cqs (see section 1, result 5), the effect of more modest non-zero values of *ρ* is slight. For example, figure S1 shows how repeatability of *mTS* relates to the error ρ values under three different assumptions about *ρ*, namely zero correlation (repeating figure 4A of the main paper), a weak correlation, and a strong correlation. Even assuming a strong correlation, error σ values of less than around 0.15 are still necessary for repeatabilities above 0.6.

**Figure S1.**
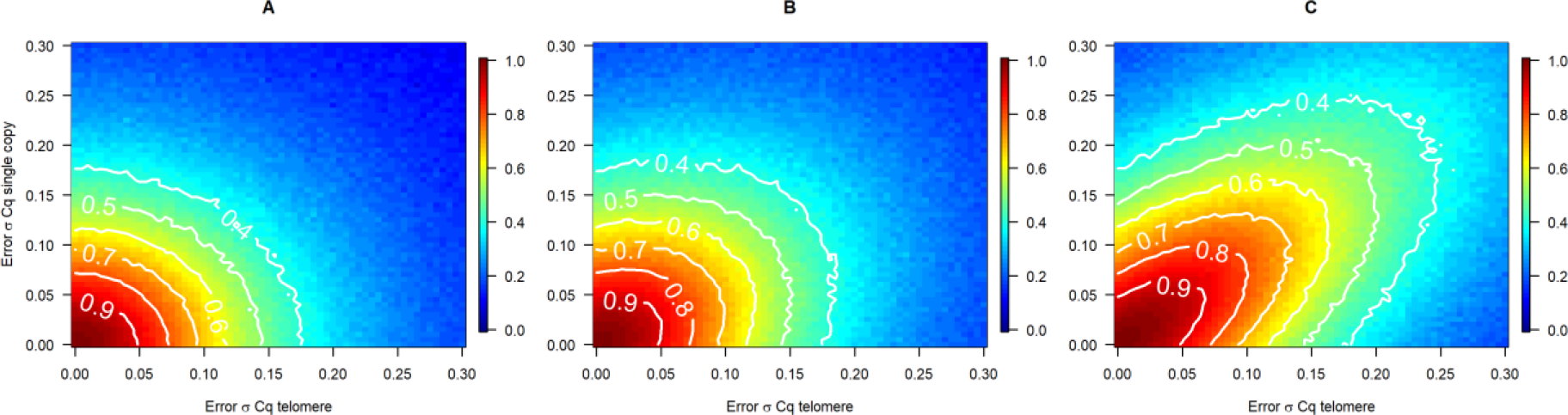
Repeatability of the TS ratio (intra-class correlation coefficient) as the error σ values for the telomere assay and the single copy gene vary. A: Errors are uncorrelated. B: Weak positive correlation (*ρ* = 0.3) between the errors. C: Strong positive correlation (*ρ* = 0.07) between the errors. Simulations of n = 10000 are used at each 0.005 step of error σ, with other parameters having their default values.

Repeating other simulated results with positive values of *ρ* produces similar conclusions: increasing *ρ* attenuates the impact of error in measuring Cqs on the TS ratio, but the effect is slight until *ρ* is close to 1.

## Section 3: How to use the simulation R code

We define a series of R functions, contained in the script ‘simulation.functions.r’, that return datasets with requested properties containing both the true values of the quantities (Cqs, TS, etc.), and their post-error measured values. This allows the user to determine the differences between true and measured values, and perform other analyses. All simulation parameter values are user-specifiable. The script ‘paper.results.r’ reproduces all the figures and simulation results from the main paper.

Datasets consist of observations from *n* individuals. The steps common to all of the simulation functions are as follows:

- A vector of *n* true single copy gene abundances, *true.dna.scg* is defined, drawn from a normal distribution with mean *b* and standard deviation *var.sample.size* (*b* is a constant).
- A vector of *n* relative telomere lengths, *true.telo.var* is defined, drawn from a normal distribution with mean 1 and standard deviation *telomere.var*.
- Hence, the true abundance of the telomere sequence is defined, as *a*true.dna.scg*true.telo.var*. Here, *a* is a scaling constant representing how many copies of the telomeric sequence there are per single copy gene in the average sample.
- Ideal Cq values for both reactions are defined as *f − log_2_(true.dna.scg)* and *f − log_2_(true.dna.telo)*, where *f* is a constant representing the chosen fluorescence threshold.
- Measurement errors in the Cqs are generated from a normal distribution with mean 0; standard deviations given by *error.scg* and *error.telo*; and a correlation between *error.scg* and *error.telo* given by *error.cor*.
- Hence, measured Cqs are generated, which can be compared to the ideal Cq values.
- TS ratios are calculated both on the measured Cqs, and the ideal ones.

The following functions are available. Specify desired parameter values in the parenthesis, e.g. *generate.one.dataset*(*n=10000, error.telo=0.1, error.scg=0.1, error.cor=0*). Default values in the simulation functions are generally those given in table 1 of the main paper.

- *generate.one.dataset()* returns a simple dataset (one telomere measurement per individual) for chosen values of all the variables described in section 1. As well as ideal and measured Cqs, it returns ideal and measured TS ratios. It also returns the difference between the ideal and measured TS ratio, calculated two ways, computed (*error.computed*), and using equation (11) of online supplement 1 (*error.analytic*). Both methods produce the same number. This was included as an additional check of correctness of the simulation.
- *generate.repeated.measure()* returns a dataset where telomere lengths from the same individuals are measured twice, via two independent biological samples, and the true telomere length of each individual is assumed not to have changed at all. The data frame it returns is as for *generate.one.dataset()*, except that there are two of each variable (e.g. *true.ts.1, true.ts.2, measured.ts.1, measured.ts.2*, etc.).
- *calculate.repeatability()* calculates the repeatability of the measured T/S ratio (intra-class correlation coefficient) when *generate.repeated.measure()* is implemented using the given values for all the parameters. It requires prior installation of R package ‘irr’.
- *compare.repeatability()* returns the repeatability of the T/S ratio and the repeatability calculated on the raw Cq for the telomere reaction, for the given parameter values.

